# Single Individual Haplotype Reconstruction Using Fuzzy C-Means Clustering With Minimum Error Correction

**DOI:** 10.1101/2020.10.21.348607

**Authors:** Mohammad Hossein Olyaee, Alireza Khanteymoori

## Abstract

Evolution of human genetics is one of the most interesting areas for researchers. Determination of Haplotypes not only makes valuable information for this purpose but also performs a major role in investigating the probable relation between diseases and genomes. Determining haplotypes by experimental methods is a time-consuming and expensive task. Recent progress in high throughput sequencing allows researchers to use computational methods for this purpose. Although, several algorithms have been proposed but they are less accurate when the error rate of input fragments increases. In this paper, first, a fuzzy conflict graph is constructed based on the similarities of all input fragments and next, the cluster centers are used as initial centers by fuzzy c-means (FCM) algorithm. The proposed method has been tested on several real datasets and compared with some current methods. The comparison with the existing approaches shows that our method can be a complementary role among the others.

## 1. Introduction

The sequencing efforts of Human genome project revealed that more than 99% of DNA sequences of human are identical [1]. As a result, the genomic differences is the responsible for diversities in our phenotypes and can be considered for many applications such as medical, drug designing, disease diagnosis and studying population history [2, 3]. Single Nucleotide Polymorphisms (SNPs) are the sites on DNA sequences that have common variations [4]. The nucleotides involved in an SNP are called alleles. Haplotype is a set of the number of SNPs that are located in a specific chromosome. Recent works show that haplotypes have more valuable information than individual SNPs [5]. In diploid organisms, such as humans, genomes are organized into pairs of chromosomes one inherited from father and other inherited from mother that are called paternal and maternal respectively. Consequently, from each copy one haplotype sequence can be gained [6, 7]. Determination of haplotypes from experimental works is so time-consuming and costing. Hence, using of computational methods is appropriate. In order to solve haplotype reconstruction problem, various methods have been proposed. At present, there are two chief models: haplotype inference [8–13] and haplotype assembly [6, 14–20]. The presented method in this article is based on the haplotype assembly.

Lancia and his colleagues[21] first proposed haplotype assembly problem. Suppose there are some short SNP fragments that are bellowing to a pair of chromosomes. Their model tries to divide these fragments in to two clusters such that each haplotype is reconstructed. Existence of errors in fragments and gaps as well as diploid organism lead to this problem becomes challenging and more difficult. Due to finding and rectifying fragment’s errors, several models have been proposed which Minimum Fragment Removal (MFR), Minimum SNP Removal (MSR), Longest Haplotype Reconstruction (LHR) and Minimum Error Correction (MEC) are four main chief models. MEC has been presented by Lippert and coworkers [22]. Although, this model is the most complicated amongst the others, it is so popular and has been used in many related works. It is proved that MEC problem is NP-hard [23].

Up to now, several approaches have been proposed to address the SIH problem based on MEC model which can be categorized as exact, metaheuristic and probabilistic methods. Exact based methods attempt to address the problem accurately and reconstruct haplotypes optimally. However, these approaches has to contain some constraints for input fragments [18, 24–26]. Since MEC problem is NP-hard, metaheuristic algorithms such as GA and PSO have been applied to solve this problem. In this case, the objective function has been designed based on MEC model and the method attempts to enhance it iteratively [6, 15, 27–30]. Existing gaps and errors in the input data, encouraged some researchers to propose probabilistic models to solve this problem. For example, HASH[19] and CUT[20] are two main approaches which lie in this category.

Fasthap method was recently proposed by Mazrouee and her colleagues [31]. Developing in accuracy and time complexity are the main goals of their method. Algorithmically, dissimilarity of every pair of fragments is measured by a new distance metric; next, a weighted graph is built based on the obtained measures; then, the created graph is used to partitioning the fragments one after another; eventually, the initial partitioning is developed in order to improve the overall MEC. The experimental results not only outperform but also the time complexity is enhanced.

Fuzzy c-means (FCM) clustering is an unsupervised technique that has been widely applied in many fields such as geology, medical imaging, target recognition, and image segmentation [32–37]. The main advantage of this approach against hard c-means is that each sample can belong to several clusters based on the measure of its membership degree. This ability is more suitable in concerning with noisy data and decrease its sensitivity against the existing noise [38]. Single individual haplotype (SIH) reconstruction problem is one of the active research areas in bioinformatics which can be modelled as a clustering problem. Most of the existing methods cluster the input fragments based on their distances. However, existing errors and gaps in the input fragments lead to computing their distances becomes unreliable.

This paper focuses on using FCM approach and introduces a new haplotype reconstruction method. Membership degree of each input fragment can interpret their belongings more precisely. The proposed method includes two steps. First, the suggested distance metric in[31] is used for building fuzzy conflict graph. Then the input fragments are partitioned into two clusters based on their similarities and from each cluster an initial haplotype is gained. In the next step, the obtained haplotypes are used by fuzzy c-means (FCM) clustering method as preliminary centers and tries to improve the accuracy of the pervious partitioning. From the results of several experiments on real data, we can see that the proposed method can always find good solutions. Also, comparing the results with several methods indicates that our method achieves an appropriate accuracy in most cases.

The rest of this paper is organized as follows: In section 2, SIH problem is formally defined and preliminary definitions and notations are given. In the third section data representation is discussed and the proposed method is described. In section 4, the experimental results of comparing the method with other popular approaches are provided. Our conclusions are drawn in the final section.

## 2. Problem formulation

Given a set of SNP fragments which are read from both chromosomes and the columns with identical values have been removed. Next as can be seen in Fig. 1 (a), an *m* × *n* matrix called SNP matrix is constructed which contains the fragments where *m* is the number of fragments and *n* is the number of the sites. In reality, there are two possible alleles for each SNP. Therefore, alleles of each SNP based on their frequency in population can be denoted by ‘*0*’ and ‘*1*’ [7] (Fig. 1 (b)).

**Fig. 1.**
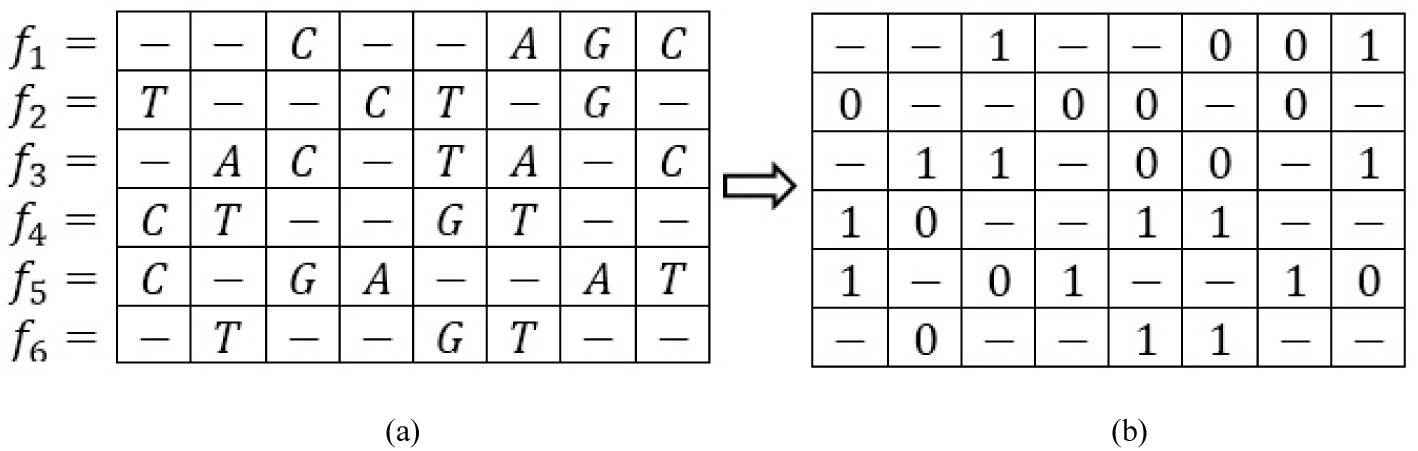
Example of SNP fragments. (a) An SNP matrix with original measures, (b) The SNP matrix which its elements have been transformed to 0/1.

Each element of matrix can be *1, 0* or ‘− ‘where ‘− ‘indicates a gap. The original haplotypes are a pair of binary strings *H(h*_*1*_,*h*_*2*_) with length *n*. The aim of SIH reconstruction is division the SNP matrix into two parts by row, and then the corresponding haplotype from each part is reconstructed.

If the fragments are error-free (Fig. 1) then they can be clustered into two groups such that all the fragments in each cluster are compatible (Fig. 2). However, in the presence of errors, there are some fragments which have conflict with the both clusters. In this case, we should reconstruct the haplotypes such that some objective function is minimized. In this study, we define this function based on Minimum error correction (MEC)[21, 22].

**Fig. 2.**
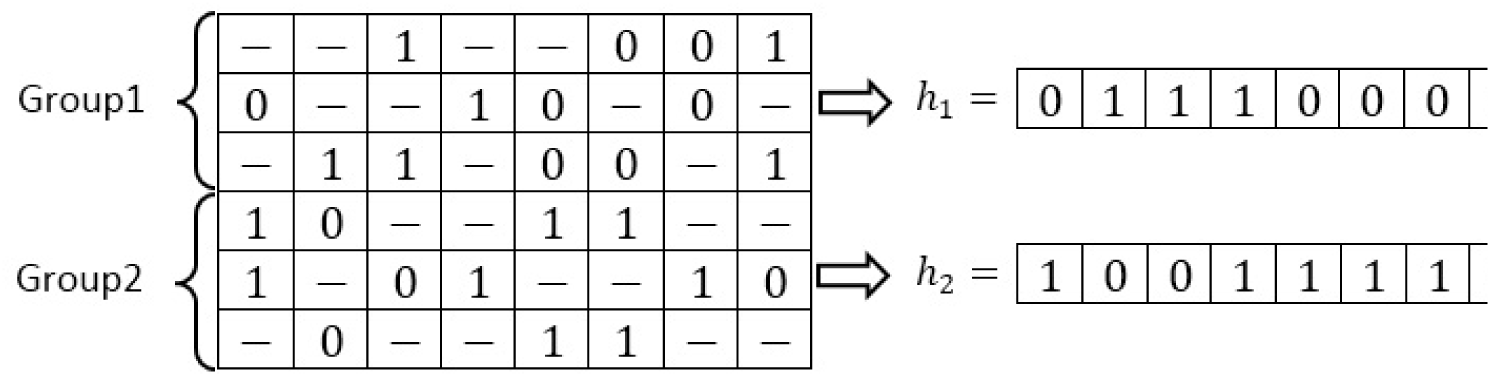
Input fragments have been divided into two groups based on their similarities and *h*_l_ and *h*_2_ are reconstructed from each group individually.

Suppose that 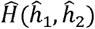 is the pair of reconstructed haplotypes. The accuracy of the algorithm is measured by reconstruction rate (RR)[6] which is defined as follows:

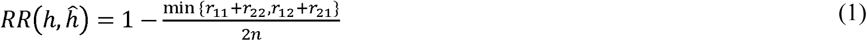

Where 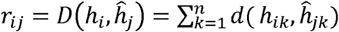 and 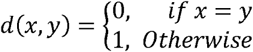

## 3. Materials and methods

As demonstrated by a series of recent publications[14, 39–46] and summarized as Chou’s 5-step rule[47], to present a suitable analysis method for a biological system, we should follow the following five guidelines: (a) select a valid benchmark dataset; (b) formulate data with an effective mathematical expression; (c) introduce a powerful algorithm to operate the reconstruction; (d) evaluate the accuracy; (e) establish a user-friendly web-server. Below, we are to describe how to deal with these steps one-by-one.

### 3.1. Materials

The Geraci’s dataset[48] is one of the major benchmarks which is prepared based on Hapmap project. This dataset consists of 22 pairs of human chromosomes from four different populations and has widely been used by several researchers [14, 15, 17, 48–50]. There are three parameters related to the data set: haplotype length, error rate and coverage rate which are denoted by *l, e, c*, respectively. Each parameter has several different values, *l* = *100, 350*, and *700, e* = *0*.*0, 0*.*1, 0*.*2* and *0*.*3, c* = *3, 5, 8* and *10*. Error rate refers to the amount of read data which has been read imprecisely. For example when e equals 0.1, it means that 10% of available data is noisy. Moreover, the coverage parameter refers to the number of times each of the two haplotypes replicates when generating the dataset. For each combination of these parameters there are *100* instances.

### 3.2. Data formulation

As it is mentioned in the previous section, suppose input SNP fragments as a *m* × *n* matrix called SNP matrix. The similarity between each two fragment *X* = (*x*_1_,*x*_2_, … *x*_*n*_) and *Y* = (*y*_1_,*y*_2_, … *y*_*n*_) can be defined as follows:

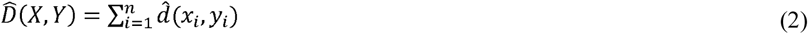

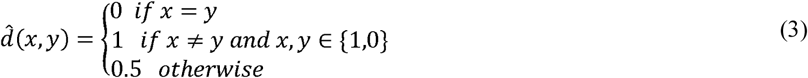

It is be noted that Eq.3 is a type of Hamming distance which has been used in [31]. It is required that the distances between all the input fragments are calculated. Next, a complete fuzzy conflict graph is constructed which has *m* vertices equal to the number of fragments and each edge representing the distance between two corresponding fragments. In fact, this graph represents dissimilarity between pairs of fragments. For example, the demonstrated matrix in Fig. 3 represents the normalized distances between all six fragments in the Fig. 1. It is be noted that distance between *f*_*i*_ and *f*_*j*_ is normalized by the number of SNP sites which at least have been covered by *f*_*i*_ or *f*_*j*_. Moreover, its corresponding fuzzy conflict graph is depicted too.

**Fig. 3.**
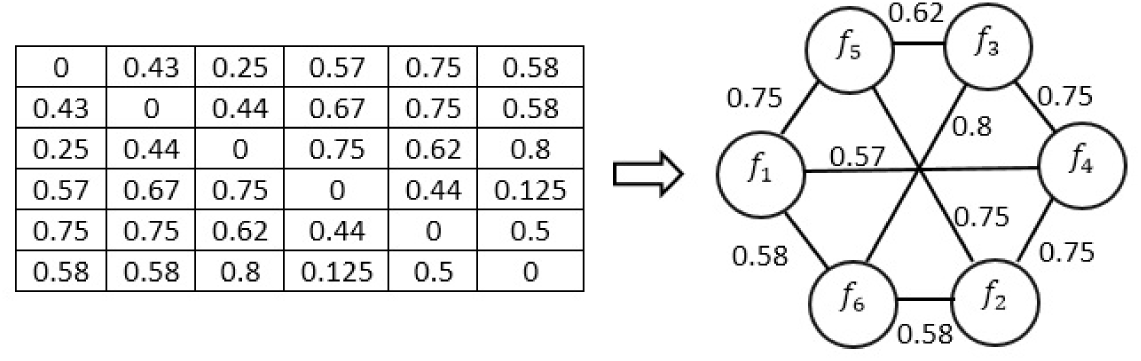
Distance matrix and corresponding fuzzy conflict graph for six input fragments

### 3.3. Proposed method

The proposed method has two phases. First, distances between all the fragments are calculated according to Eq.3 and the corresponding fuzzy conflict graph is constructed. Next, the obtained distances are used to bi-partitioning all the fragments. It is be noticed that this clustering is done based on the dissimilarities between the fragments. In the second phase, centers of the gained clusters (the obtained haplotypes) are used as initial centers by fuzzy c-means algorithm. First step leads to increase the convergence speed of FCM and decreases the number of iterations. The FCM algorithm assigns fragments to each cluster according to fuzzy memberships. Let 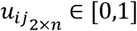 is degree of membership of *jth* fragment to *ith* cluster which *i* ∈ {0,1}and matrix U = [*u*_*ij*_]_2 × *n*_ contains the membership of all the fragments. In this way each fragment may belong to any of the two clusters by different membership degrees. The algorithm is an iterative optimization that minimizes the cost function defined as bellows:

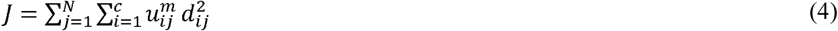

Where *d*_*ij*_ the distance between each cluster center and input fragments that defined is based on relation (1), N is the number of fragments and m is a constant which controls the fuzziness of the resulting partition. This measure can be set between one to infinity and there isn’t any theoretical way to determine it. In this study, m equals with 2 based on several past researches [33, 37, 51–53]. The updated membership matrix and cluster centers are calculated from

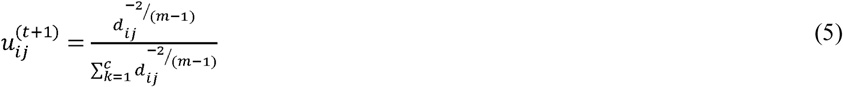

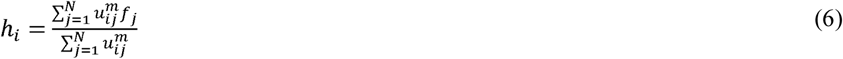

The algorithm is expressed by the flowchart shown in Fig. 4. The last two steps are iterated until the improvement over the previous iteration is below a threshold *ε*. The cost function is minimized when fragments close to the centroid of their clusters are assigned high membership values, and low membership values are assigned to fragments that far from the centroid.

**Fig. 4.**
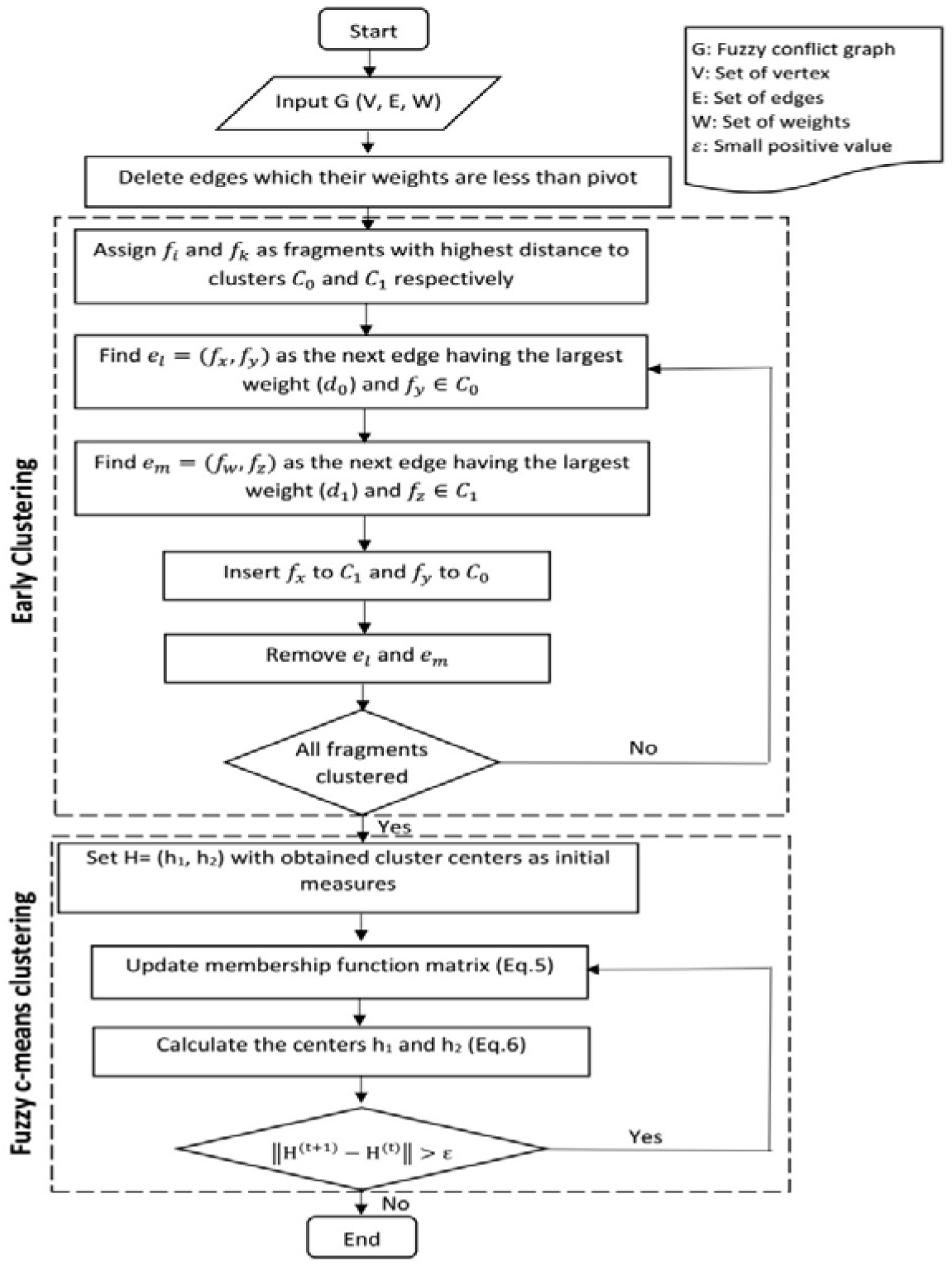
Flowchart of the proposed method

## 4. Experimental results

To evaluate the performance of our method, as mentioned previously, we have used the dataset in Geraci’s research [48, 54].

In order to assess the performance of the proposed algorithm, it is compared with the algorithms that were investigated in Geraci’s research[48]. MLF[55] and 2d [26] are based on MEC model. The former uses confidence score for each SNP site and the later uses two distance metrics in order to cluster input fragments. Both of DGS[56] and Cut[20] methods works with a sub-matrix of input fragments. The first, considers a pair of haplotypes as initial and based on the majority rule refines it step by step. The second models the SIH as a max-cut problem in derived SNP graphs. Fast[57] sorts input fragments according to the positions which gaps begin and assigns them to the clusters. SHR[58] is a randomized-based approach which selects the input fragments in an iterative manner and by exploiting hamming distance assigns them to the closest set. Finally, SPH[54] is a heuristic-based method expoliting the statistical correlations between SNPs and uses for the high noisy fragments with low coverages. The results of proposed method called FCMhap as well as the results of other algorithms can be seen as follows. It should be noticed that Table 1–3 represents the results of haplotypes with length *100, 350* and *700* respectively. The first two columns in these tables indicate error rate and coverage separately. The results of our method can be seen in the last column. The bold values specify the utmost RRs, also the gray values indicate the second highest RRs. It is interesting to note that all compering methods have good performance in the error-free cases. However, by increasing the amount of noise and gaps, their performances decrease dramatically. By using FCM, each fragment can be belong to both clusters. Their belonging have been determined according to their membership degree measures. The membership degree can describe the belonging of each fragment more accurately especially for input fragments with large amount of noise. Therefore, as can be seen here, the comparison of results particularly in cases with a high error rate, demonstrates that FCMhap has suitable performance against the other methods.

**Table 1.** The average of reconstruction rate for 100 examples with length 100

| e | c | SPH | Fast | 2d | Cut | MLF | SHR | DGS | FCMhap |
| --- | --- | --- | --- | --- | --- | --- | --- | --- | --- |
| **0** | 3 | 0.999 | 0.999 | 0.990 | 1.000 | 0.973 | 0.816 | 1.000 | **1.000** |
| **0** | 5 | 1.000 | 0.999 | 0.997 | 1.000 | 0.992 | 0.861 | 1.000 | **1.000** |
| **0** | 8 | 1.000 | 1.000 | 1.000 | 1.000 | 0.997 | 0.912 | 1.000 | **1.000** |
| **0** | 10 | 1.000 | 1.000 | 1.000 | 1.000 | 0.998 | 0.944 | 1.000 | **1.000** |
| **0.1** | 3 | 0.895 | 0.913 | 0.911 | 0.928 | 0.889 | 0.696 | 0.930 | 0.8816 |
| **0.1** | 5 | 0.967 | 0.964 | 0.951 | 0.920 | 0.969 | 0.738 | 0.985 | 0.9478 |
| **0.1** | 8 | 0.989 | 0.993 | 0.983 | 0.901 | 0.985 | 0.758 | 0.989 | 0.9713 |
| **0.1** | 10 | 0.990 | 0.998 | 0.988 | 0.892 | 0.995 | 0.762 | 0.997 | 0.9725 |
| **0.2** | 3 | 0.623 | 0.715 | 0.738 | 0.782 | 0.725 | 0.615 | 0.725 | 0.7392 |
| **0.2** | 5 | 0.799 | 0.797 | 0.793 | 0.838 | 0.836 | 0.655 | 0.813 | 0.7721 |
| **0.2** | 8 | 0.852 | 0.881 | 0.873 | 0.864 | 0.918 | 0.681 | 0.878 | 0.7926 |
| **0.2** | 10 | 0.865 | 0.915 | 0.894 | 0.871 | 0.938 | 0.699 | 0.917 | 0.8348 |
| **0.3** | 3 | 0.480 | 0.617 | 0.623 | 0.602 | 0.618 | 0.557 | 0.611 | **0.6286** |
| **0.3** | 5 | 0.637 | 0.639 | 0.640 | 0.629 | 0.653 | 0.599 | 0.647 | 0.6485 |
| **0.3** | 8 | 0.667 | 0.661 | 0.675 | 0.673 | 0.697 | 0.632 | 0.663 | 0.6643 |
| **0.3** | 10 | 0.676 | 0.675 | 0.678 | 0.709 | 0.715 | 0.632 | 0.688 | 0.6754 |

**Table 2.** The average of reconstruction rate for 100 examples with length 350

| e | c | SPH | Fast | 2d | Cut | MLF | SHR | DGS | FCMhap |
| --- | --- | --- | --- | --- | --- | --- | --- | --- | --- |
| **0** | 3 | 0.999 | 0.989 | 0.965 | 1.000 | 0.864 | 0.830 | 1.000 | **1.000** |
| **0** | 5 | 1.000 | 0.999 | 0.993 | 1.000 | 0.929 | 0.829 | 1.000 | **1.000** |
| **0** | 8 | 1.000 | 1.000 | 0.998 | 1.000 | 0.969 | 0.895 | 1.000 | **1.000** |
| **0** | 10 | 1.000 | 1.000 | 0.999 | 1.000 | 0.981 | 0.878 | 1.000 | **1.000** |
| **0.1** | 3 | 0.819 | 0.871 | 0.839 | 0.930 | 0.752 | 0.682 | 0.926 | 0.873 |
| **0.1** | 5 | 0.959 | 0.945 | 0.913 | 0.913 | 0.858 | 0.7244 | 0.978 | 0.9186 |
| **0.1** | 8 | 0.984 | 0.985 | 0.964 | 0.896 | 0.933 | 0.742 | 0.996 | 0.9344 |
| **0.1** | 10 | 0.984 | 0.995 | 0.978 | 0.888 | 0.962 | 0.728 | 0.998 | 0.935 |
| **0.2** | 3 | 0.439 | 0.684 | 0.675 | 0.771 | 0.642 | 0.591 | 0.691 | 0.671 |
| **0.2** | 5 | 0.729 | 0.746 | 0.728 | 0.831 | 0.728 | 0.632 | 0.769 | 0.7186 |
| **0.2** | 8 | 0.825 | 0.853 | 0.791 | 0.862 | 0.798 | 0.670 | 0.842 | 0.7279 |
| **0.2** | 10 | 0.855 | 0.877 | 0.817 | 0.867 | 0.831 | 0.668 | 0.878 | 0.7335 |
| **0.3** | 3 | 0.251 | 0.590 | 0.593 | 0.565 | 0.581 | 0.548 | 0.578 | **0.5975** |
| **0.3** | 5 | 0.578 | 0.602 | 0.606 | 0.582 | 0.606 | 0.557 | 0.609 | **0.6137** |
| **0.3** | 8 | 0.629 | 0.626 | 0.623 | 0.621 | 0.634 | 0.604 | 0.628 | 0.6264 |
| **0.3** | 10 | 0.638 | 0.644 | 0.634 | 0.664 | 0.641 | 0.619 | 0.641 | 0.631 |

**Table 3.** The average of reconstruction rate for 100 examples with length 700

| e | c | SPH | Fast | 2d | Cut | MLF | SHR | DGS | FCMhap |
| --- | --- | --- | --- | --- | --- | --- | --- | --- | --- |
| **0** | 3 | 0.999 | 0.988 | 0.946 | 1.000 | 0.782 | 0.781 | 1.000 | **1.000** |
| **0** | 5 | 1.000 | 0.999 | 0.976 | 1.000 | 0.854 | 0.832 | 1.000 | **1.000** |
| **0** | 8 | 1.000 | 1.000 | 0.992 | 1.000 | 0.919 | 0.868 | 1.000 | **1.000** |
| **0** | 10 | 1.000 | 0.999 | 0.997 | 1.000 | 0.933 | 0.898 | 1.000 | **1.000** |
| **0.1** | 3 | 0.705 | 0.829 | 0.786 | 0.927 | 0.698 | 0.668 | 0.931 | 0.8344 |
| **0.1** | 5 | 0.947 | 0.941 | 0.880 | 0.916 | 0.809 | 0.716 | 0.977 | 0.881 |
| **0.1** | 8 | 0.985 | 0.986 | 0.948 | 0.896 | 0.863 | 0.743 | 0.987 | 0.8833 |
| **0.1** | 10 | 0.986 | 0.995 | 0.965 | 0.889 | 0.884 | 0.726 | 0.997 | 0.996 |
| **0.2** | 3 | 0.199 | 0.652 | 0.647 | 0.753 | 0.624 | 0.591 | 0.669 | 0.6517 |
| **0.2** | 5 | 0.681 | 0.712 | 0.697 | 0.825 | 0.682 | 0.617 | 0.741 | 0.6718 |
| **0.2** | 8 | 0.801 | 0.808 | 0.751 | 0.856 | 0.747 | 0.653 | 0.818 | 0.6863 |
| **0.2** | 10 | 0.813 | 0.872 | 0.778 | 0.861 | 0.765 | 0.675 | 0.861 | 0.7458 |
| **0.3** | 3 | 0.095 | 0.581 | 0.583 | 0.552 | 0.570 | 0.536 | 0.573 | **0.5923** |
| **0.3** | 5 | 0.523 | 0.591 | 0.596 | 0.555 | 0.594 | 0.562 | 0.595 | **0.5988** |
| **0.3** | 8 | 0.616 | 0.615 | 0.613 | 0.597 | 0.614 | 0.611 | 0.614 | 0.606 |
| **0.3** | 10 | 0.627 | 0.616 | 0.622 | 0.645 | 0.625 | 0.625 | 0.622 | 0.6064 |

In this study, we have focused on the improvement of the reconstruction rate. However, in order to provide a comprehensive assessment about the proposed method, its running time has been compared against the other approaches. For this purpose, for each combination of the parameters, the methods have been run by an ordinary desktop PC over 10 samples which have been selected randomly. The average of running times for each set of parameters have been collected which can be seen in Tables 4–6.

**Table 4.** The average of runing time for examples with length 100 (in seconds)

| e | c | SPH | Fast | 2d | Cut | MLF | SHR | DGS | FCMhap |
| --- | --- | --- | --- | --- | --- | --- | --- | --- | --- |
| 0 | 3 | 0.006 | 0.001 | 0.004 | 0.013 | 0.012 | 0.002 | 0.001 | 0.018 |
| 0 | 5 | 0.009 | 0.001 | 0.003 | 0.022 | 0.029 | 0.001 | 0.002 | 0.026 |
| 0 | 8 | 0.013 | 0.002 | 0.013 | 0.033 | 0.046 | 0.002 | 0.006 | 0.055 |
| 0 | 10 | 0.016 | 0.003 | 0.016 | 0.136 | 0.067 | 0.003 | 0.016 | 0.047 |
| 0.1 | 3 | 0.007 | 0.001 | 0.004 | 0.060 | 0.012 | 0.001 | 0.001 | 0.017 |
| 0.1 | 5 | 0.009 | 0.002 | 0.006 | 0.083 | 0.023 | 0.001 | 0.005 | 0.027 |
| 0.1 | 8 | 0.013 | 0.002 | 0.012 | 0.107 | 0.052 | 0.002 | 0.009 | 0.039 |
| 0.1 | 10 | 0.014 | 0.003 | 0.024 | 0.162 | 0.071 | 0.003 | 0.011 | 0.051 |
| 0.2 | 3 | 0.007 | 0.001 | 0.004 | 0.053 | 0.013 | 0.001 | 0.001 | 0.018 |
| 0.2 | 5 | 0.011 | 0.001 | 0.008 | 0.085 | 0.019 | 0.001 | 0.005 | 0.027 |
| 0.2 | 8 | 0.008 | 0.002 | 0.017 | 0.146 | 0.040 | 0.003 | 0.010 | 0.041 |
| 0.2 | 10 | 0.014 | 0.002 | 0.016 | 0.185 | 0.050 | 0.004 | 0.009 | 0.046 |
| 0.3 | 3 | 0.008 | 0.001 | 0.003 | 0.093 | 0.009 | 0.001 | 0.001 | 0.015 |
| 0.3 | 5 | 0.011 | 0.001 | 0.007 | 0.093 | 0.028 | 0.002 | 0.004 | 0.029 |
| 0.3 | 8 | 0.011 | 0.002 | 0.014 | 0.176 | 0.038 | 0.002 | 0.009 | 0.038 |
| 0.3 | 10 | 0.013 | 0.002 | 0.022 | 0.182 | 0.071 | 0.002 | 0.017 | 0.044 |

**Table 5.** The average of running time for examples with length 350 (in seconds)

| **e** | **c** | **SPH** | **Fast** | **2d** | **Cut** | **MLF** | **SHR** | **DGS** | **FCMhap** |
| --- | --- | --- | --- | --- | --- | --- | --- | --- | --- |
| 0 | 3 | 0.212 | 0.009 | 0.055 | 0.178 | 0.197 | 0.010 | 0.047 | 0.196 |
| 0 | 5 | 0.369 | 0.011 | 0.166 | 0.352 | 0.457 | 0.013 | 0.107 | 0.273 |
| 0 | 8 | 0.363 | 0.016 | 0.453 | 0.385 | 0.533 | 0.027 | 0.212 | 0.493 |
| 0 | 10 | 0.428 | 0.031 | 0.756 | 0.657 | 1.384 | 0.032 | 0.329 | 0.604 |
| 0.1 | 3 | 0.281 | 0.008 | 0.050 | 2.990 | 0.215 | 0.007 | 0.034 | 0.162 |
| 0.1 | 5 | 0.406 | 0.017 | 0.208 | 4.762 | 0.263 | 0.017 | 0.106 | 0.291 |
| 0.1 | 8 | 0.398 | 0.015 | 0.504 | 10.140 | 0.525 | 0.026 | 0.264 | 0.469 |
| 0.1 | 10 | 0.431 | 0.019 | 0.625 | 15.785 | 1.316 | 0.032 | 0.271 | 0.621 |
| 0.2 | 3 | 0.231 | 0.009 | 0.065 | 3.985 | 0.155 | 0.010 | 0.032 | 0.185 |
| 0.2 | 5 | 0.296 | 0.017 | 0.129 | 8.534 | 0.445 | 0.011 | 0.110 | 0.284 |
| 0.2 | 8 | 0.443 | 0.018 | 0.346 | 12.543 | 0.566 | 0.015 | 0.269 | 0.509 |
| 0.2 | 10 | 0.495 | 0.029 | 0.718 | 18.354 | 1.423 | 0.029 | 0.408 | 0.596 |
| 0.3 | 3 | 0.380 | 0.009 | 0.072 | 4.734 | 0.196 | 0.009 | 0.030 | 0.163 |
| 0.3 | 5 | 0.400 | 0.015 | 0.188 | 6.252 | 0.458 | 0.015 | 0.081 | 0.276 |
| 0.3 | 8 | 0.526 | 0.015 | 0.404 | 15.668 | 0.712 | 0.017 | 0.242 | 0.455 |
| 0.3 | 10 | 0.486 | 0.024 | 0.736 | 16.343 | 1.881 | 0.022 | 0.460 | 0.679 |

**Table 6.** The average of running time for examples with length 700 (in seconds)

| **e** | **c** | **SPH** | **Fast** | **2d** | **Cut** | **MLF** | **SHR** | **DGS** | **FCMhap** |
| --- | --- | --- | --- | --- | --- | --- | --- | --- | --- |
| 0 | 3 | 4.328 | 0.022 | 0.508 | 0.855 | 1.235 | 0.028 | 0.352 | 1.601 |
| 0 | 5 | 4.043 | 0.059 | 2.009 | 1.513 | 2.689 | 0.037 | 0.650 | 3.025 |
| 0 | 8 | 3.935 | 0.064 | 5.537 | 3.696 | 4.371 | 0.100 | 3.581 | 4.381 |
| 0 | 10 | 5.897 | 0.134 | 6.460 | 3.825 | 9.093 | 0.124 | 6.084 | 6.866 |
| 0.1 | 3 | 3.060 | 0.050 | 0.822 | 38.525 | 2.029 | 0.051 | 0.351 | 1.613 |
| 0.1 | 5 | 2.781 | 0.035 | 2.110 | 40.845 | 2.477 | 0.060 | 0.614 | 2.580 |
| 0.1 | 8 | 3.588 | 0.076 | 4.937 | 82.105 | 4.546 | 0.058 | 2.308 | 4.226 |
| 0.1 | 10 | 5.743 | 0.162 | 7.153 | 134.104 | 8.764 | 0.136 | 5.989 | 6.765 |
| 0.2 | 3 | 3.310 | 0.033 | 0.333 | 24.271 | 0.757 | 0.026 | 0.400 | 1.609 |
| 0.2 | 5 | 3.529 | 0.047 | 2.132 | 54.829 | 2.421 | 0.055 | 0.970 | 2.524 |
| 0.2 | 8 | 4.408 | 0.084 | 5.141 | 102.456 | 6.004 | 0.067 | 4.169 | 4.558 |
| 0.2 | 10 | 5.274 | 0.102 | 7.826 | 100.052 | 6.737 | 0.110 | 5.118 | 6.410 |
| 0.3 | 3 | 2.755 | 0.021 | 0.332 | 41.529 | 0.713 | 0.026 | 0.306 | 1.612 |
| 0.3 | 5 | 3.867 | 0.061 | 1.674 | 63.359 | 1.439 | 0.063 | 0.965 | 2.874 |
| 0.3 | 8 | 3.912 | 0.097 | 5.368 | 119.681 | 3.764 | 0.070 | 3.464 | 4.918 |
| 0.3 | 10 | 5.403 | 0.111 | 8.848 | 173.903 | 7.265 | 0.108 | 6.828 | 6.983 |

The comparison of the haplotype reconstruction time demonstrates that the running time of the proposed method is scalable with the other approaches and it can reconstruct haplotypes for each parameter assignment in less than seven seconds.

## 5. Conclusion

Providing huge amount of genomic sequences has been increased the importance of single individual haplotype problem. Determination of haplotype can be useful in several domains such as understanding the relation between genetic variations and complicated diseases. Since laboratory-based methods are time consuming and expensive, several computational-based approaches have been proposed which reconstruct haplotypes directly from the reads. But their performance can be dramatically decreased in dealing with the noisy input data. We have presented FCMhap, an effective method that utilizes Fuzzy c-means (FCM) algorithm as a main step. FCM by considering a fuzzy membership for each read can efficiently cluster the noisy data. The obtained results demonstrate that FCMhap can improve reconstruction rate especially for high-error-rate data. It should be noted that the codes used to prepare this article are available from the author upon request.

## Acknowledgement

We would like to acknowledge the help that received from our colleagues in Machine Learning and Bioinformatics Laboratory (MLBL) of University of Zanjan, Zanjan, Iran. The authors would also like to thank Dr. F. Geraci for providing his benchmark data set.

## Notes

### Competing Interest Statement

The authors have declared no competing interest.

